# Coordinated shifts in allocation among reproductive tissues across 14 coexisting plant species

**DOI:** 10.1101/141473

**Authors:** E. H. Wenk, K. Abramowicz, M. Westoby, D. S. Falster

**Affiliations:** Department of Biological Sciences, Macquarie University NSW 2109, Australia; Department of Mathematics and Mathematical Statistics, Umeå University, 90187 Umeå, Sweden; Evolution and Ecology Research Centre, School of Biological, Earth and Environmental Science, University of New South Wales

**Keywords:** reproduction, accessory costs, parental optimist, seed provisioning, selective abortion, seed size-number trade-off

## Abstract

Plant species differ in the amounts of energy allocated to different reproductive tissues, driving differences in their ecology and energy flows within ecosystems. While it is widely agreed that energy allocation is key to reproductive outcomes, few studies have estimated how reproductive effort (RE) is partitioned among different pools, for multiple species in a community. In plants, RE can be partitioned in several meaningful ways: seed versus non-seed tissues; into flowers that form seeds and those that fail to develop; into pre-versus post-pollination tissues, and into successful versus aborted ovules. Evolutionary theory suggests several hypotheses about how these tissues should be coordinated across species. To quantify variation in allocation to different reproductive tissues, we collected detailed RE measurements for a year from 14 perennial species in a recurrent-fire coastal heath community in eastern Australia. Overall we found that total accessory costs – the proportion of RE not directly invested in provisioning the seed – were very large, varying from 95.8% to 99.8% across the study species. These results suggest that studies using seed or fruit production as measures of RE may underestimate it by 10-to 500-fold. We propose a suitable alternative that well-approximates true RE. When comparing species, we found strong support for three evolutionary trade-offs that are predicted to arise when a given energy pool is divided into different tissue masses and counts across species: 1) between successful pollen-attraction costs and mature ovule count, 2) between total reproductive costs and seed count, and 3) between seedset and relative investment in pollen-attraction costs. As a result of these trade-offs, species were also predicted to show coordinated shifts in the amounts invested in floral construction, in seedset and seed size. These shifts in investment were indeed observed, with the amount allocated to discarded tissues increasing with seed size and the amount allocated to pollen-attraction decreasing with seed size. It is already well-established that the seed size axis aligns with the colonization-competition life history spectrum; here we show that relative construction costs of pollen-attraction versus provisioning tissues and seedset are also part of this trajectory, expanding our understanding of the relatives sizes of floral and fruiting structures observed across angiosperms.

## Introduction

Plants allocate a sizeable share of their photosynthetic energy to reproduction (Obeso 2004; Hirayama *et al.* 2008; Thomas 2011; Wenk & Falster 2015). This allocation takes the form of provisioned seeds and also of many other tissues associated with reproduction, termed accessory costs. Accessory costs include energy associated with forming a successful seed (e.g. flower petals, seed pod, and dispersal tissues) and energy lost via aborted and discarded buds, flowers and fruit. Previous studies show that for perennial species anywhere from 15% – 99% of total reproductive investment may go into accessory costs (Haig & Westoby 1988; Ashman 1994; Henery & Westoby 2001; Lord & Westoby 2006; Chen, Felker & Sun 2010). Since fruit set and seed set are generally below 50% in perennial species (Stephenson 1981; Wiens 1984; Sutherland 1986; Knight *et al.* 2005; Rosenheim *et al.* 2014), the cost of aborted and discarded tissues may be a substantial proportion of total accessory costs. Yet, despite being a significant energy sink in ecosystems, little is known about the allocation of energy among different reproductive tissues across the plant kingdom and how this links with plant reproductive strategies.

While plant species demonstrate an extraordinary diversity of reproductive structures and strategies, reproductive investment can be divided into broad functional categories that are consistent across species (Figure 1a). We define categories as follows. Total energy investment per seed matured is *reproductive costs*. This can be divided into investment in required parts, termed *success costs*, and energy expenditure on flowers, fruit, and seeds that never form mature propagules, called *discarded tissue costs*. All are calculated on a *per seed matured* basis. *Success costs* can be further divided into structures that form before pollination *(pollen-attraction costs;* i.e. the flower, including petals, calyx, pedicel) versus structures developed post-pollination (e.g. seed pod, seed), hereafter termed *provisioning costs*. The provisioning component is comprised of the seed itself (seed size) versus the dispersal and packaging tissues. Although in much of the literature seed mass is understandably treated as including the seed coat, for purposes of this paper we treat embryo plus endosperm mass as seed mass and position the seed coat among the dispersal and packaging component of accessory costs. The *discarded tissue costs* can likewise be divided into energy invested prior to versus after pollination, here termed *discarded pollen-attraction costs* and *discarded provisioning costs. Accessory costs*, all tissues besides the seed mass itself, are the sum of *discarded tissue costs, pollen-attraction costs, and packaging and dispersal costs*, terms high-lighted in red in Figure 1a. Throughout the manuscript “costs” indicates dry mass investment per seed matured, while “investment” refers to total dry mass invested in a structure.

**Figure 1.**
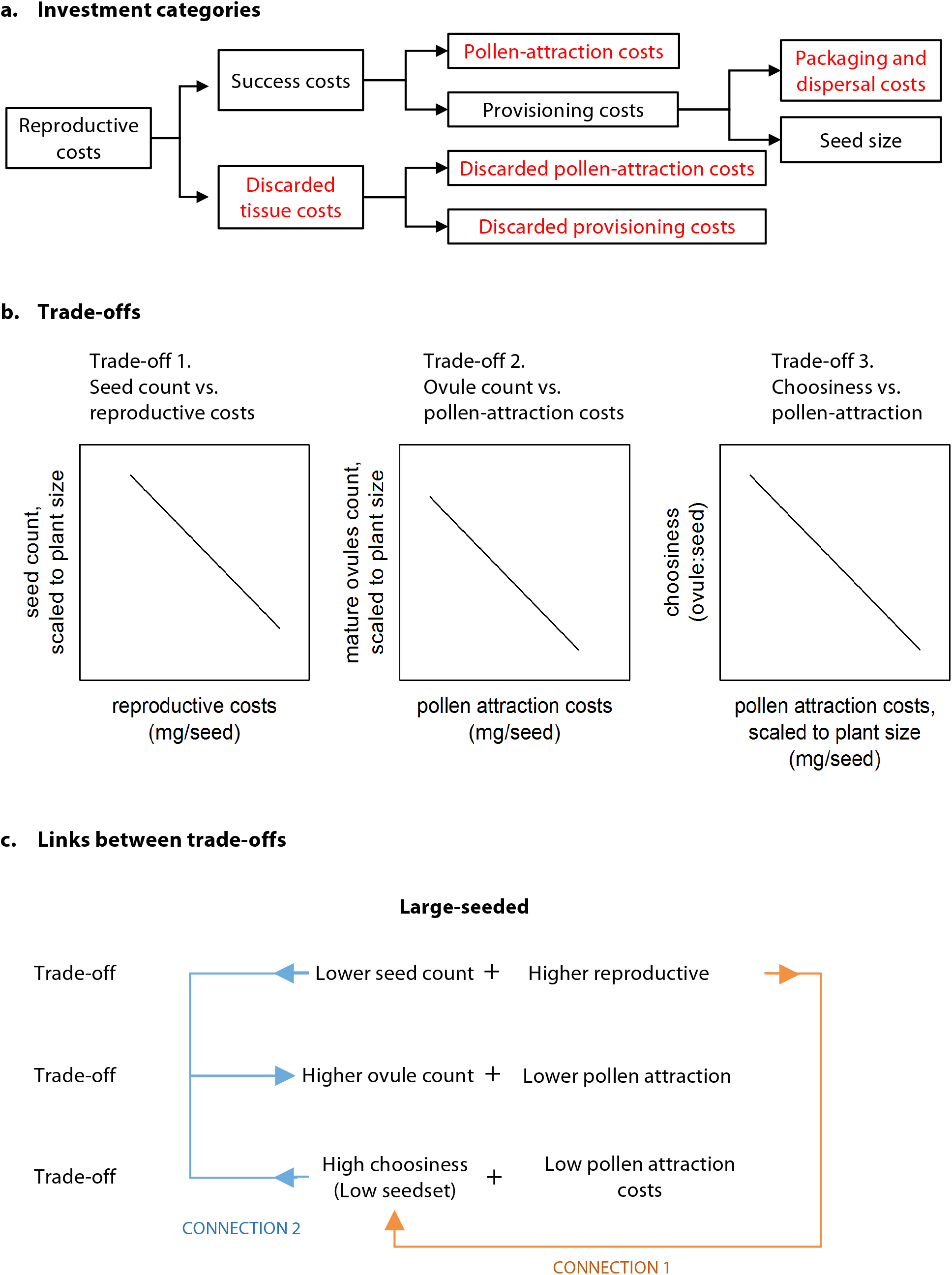
a) Categories of reproductive investment, expressed as “costs”, defined as investment divided by count of seeds matured. Categories in red are components of total accessory costs. b) Three trade-offs are predicted: Trade-off 1. For a given energy pool to be invested in total reproduction, there is a trade-off between total reproductive investment per seed produced and number of seeds produced; Trade-off 2. For a given energy pool to be invested to the point of pollination, there is a trade-off between pollen-attraction costs and the number of mature ovules produced; and Trade-off 3: A tradeoff between “choosiness”, the ratio of mature ovules to mature seeds, and pollen-attraction costs, scaled to plant size, is also predicted, for a plant with more costly pollen-attraction tissues will be able to produce fewer excess ovules. c) Together, these trade-offs predict a syndrome of reproductive traits for large versus small-seeded species, for the three trade-offs are linked by natural selection and logic. Connection 1 indicates that species with high reproductive costs will also be highly selective about which ovules to mature. Connection 2 shows that a species on the low seed-count end of trade-off 1 and the high choosiness end of trade-off 3 will, by definition, have a relatively high ovule count. The figure depicts the end of each trade-off predicted for a large-seeded species. A small-seeded species is predicted to have cost and count values at the opposite end of each trade-off.

There are multiple reasons to expect that both success costs and discarded tissue costs will be substantially larger than the mass of the seed itself. The success cost components are undeniably beneficial for successful formation and dispersal of a seed. Without showy petals insects would not be attracted to the stamens and stigma, without sepals the developing bud would not be protected, without a seed coat a seed would not be protected during dispersal, and without an attractive fruit, many seeds would not be dispersed. High discarded tissue costs (due to low seedset both pre- and post-pollination) occur in perennial plants for a diversity of reasons, some the result of conditions beyond the plant’s control and others by evolutionary design to increase fitness. They include pollen-limitation, pollen-ovule incompatibility, parental embryo abortion, resource limitation and bet-hedging strategies to capitalize on stochastic variation in pollen availability, pollen quality, and resource availability to mature fertilized ovules (Bierzychudek 1981; Stephenson 1981; Sutherland 1986; Burd 1994, 2008; Ramsey 1997; Obeso 2004; Ashman *et al.* 2004; Knight *et al.* 2005; Holland & Chamberlain 2007; Rosenheim, Schreiber & Williams 2015). Variation across species in the relative size of the reproductive tissue energy expenditures should indicate different reproductive energy allocation strategies underpinned by trade-offs (Figure 1a). Species may differ in how they divide their finite pool of reproductive energy into different tissue types, displaying variation in relative investment in pollen-attraction versus provisioning costs as well as variation in the number of ovules formed and the number of seeds matured.

The literature identifies two main reproductive strategy trade-offs relating reproductive energy pools and counts of reproductive parts to each other. Here we expand upon those hypotheses and show that they capture the same life history strategy spectrum from different perspectives. The first is the well-supported seed size-seed number trade-off, from the plant functional trait literature. The second is the seed set-pollen-attraction cost trade-off described in the parental optimist-parental pessimist literature. Each of these trade-offs is separate, for each considers different resources, yet together these yield hypotheses on how energy allocation to the energy pools illustrated in Figure 1a should differ systematically with respect to seed size through evolutionary linkages.

### Seed size – seed number trade-off

Whatever pool of energy is available to a plant for seed production can be divided into many small seeds or fewer larger seeds (Smith & Fretwell 1974). A log-log plot of seed size versus seed count scaled to plant size should have a slope of −1 all else being equal. Indeed, a slope of −1 has been observed across species in the field (Henery & Westoby 2001; Moles *et al.* 2004; Sadras 2007). Very small and very large seeds represent endpoints of a continuous spectrum of life history strategies (Rees & Westoby 1997; Leishman 2001; Turnbull *et al.* 2004; Moles & Westoby 2006). Small-seeded species have a greater chance of reaching any given colonization opportunity, while larger-seeded species have a greater likelihood of establishing and better competitive outcomes at any given location (Moles & Westoby 2006).

The seed size-seed number trade-off does not consider energy invested in accessory tissues, leading us to hypothesize two related trade-offs. First, within a given total expenditure on reproduction, there should be a trade-off between seed count and total reproductive costs per seed matured (trade-off 1 in Figure 1b). This trade-off is similar to the seed size-seed count trade-off, but includes all of a plant’s reproductive energy expenditures to construct a seed, not just the seed mass itself. Second, within a given amount of energy spent to mature ovules to the point of pollination, there should be a trade-off between pollen-attraction costs per ovule and the number of ovules that are displayed to pollinators (trade-off 2 in Figure 1b). Species with higher pollen-attraction costs are expected to produce fewer ovules. Both trade-offs are predicted to have a slope = −1, but the trade-off between pollen-attraction costs and ovules at point of pollination should have a higher intercept, since seed set per ovule is <1. These are two independent trade-offs, each simply showing there exists a fixed pool of energy to be divided among offspring. Species variation in seedset, the ratio of seed count to ovule count, provides the link between these two trade-offs, and is itself one of the axes in the trade-off described below.

### The pollen attraction-seed provisioning versus seed set trade-off

Haig & Westoby (1988) developed a conceptual model for the relative allocation of energy to different reproductive tissues, dividing the total energy investment per seed between the costs of acquiring pollen and the cost of provisioning pollinated ovules. Their simple model makes several predictions, including that plants 1) produce excess ovules and flowers to optimize seed production across a population and across time, 2) face a trade-off between pollen attraction and embryo provisioning, and 3) allocate just enough to pollen-attracting tissues to ensure pollination of the number of ovules they are able to provision *on average*. This initial model has since been extended to use the proportion of energy invested in pollen attraction versus seed provisioning tissues to predict seed set across species (Rosenheim *et al.* 2014, 2016). The models, supported by empirical data, indicate that species with relatively low pollen-attraction costs should produce a greater excess of ovules relative to what they are able to provision; in other words they should have lower seedset.

This axis of variation aligns with the parental optimist-parental pessimist strategy continuum (Mock & Forbes 1995; Burd 2008; Rosenheim *et al.* 2014). A parental optimist is a species that overproduces ovules, relatively few of which mature in an average year due to limited resource supply. Such a species is “optimistic” in the sense that should environmental conditions be unusually favorable, it will be able to respond with high seed production. Since an optimist, in average years, discards so many ovules – both pollinated and unpollinated – it must reduce the cost of producing a single pollinator-ready ovule. The alternative, a species with proportionally higher pollen-attraction costs, should display parental pessimism and produce relatively fewer ovules, with embryo number limiting seed production in many years.

Since parental optimists have lower seedset (seed to ovule ratio), logically they need to ensure that the seeds they mature are likely to germinate and establish. One mechanism to increase seed and seedling success is to invest more resources in embryo provisioning, manifested as higher packaging and dispersal costs and higher seed mass. High per seed resource investment in turn will favor provisioning embryos that are vigorous genotypes, in part accomplished by being selective about which pollen grains to use and which zygotes to provision, termed selective abortion. This has been shown to be an important mechanism to increase plant fitness (Willson & Burley 1983; Sutherland 1986; Kozlowski & Stearns 1989; Guittian 1993; Melser & Klinkhamer 2001; Harder & Barrett 2006). A parent plant can exert stronger zygote selection if a large pool of excess zygotes is brought into existence, exactly the strategy displayed by a parental optimist. In summary, we expect the ratio of ovules to seeds, defined here as *choosiness* (the inverse of *seedset*), to be highest in parental optimists, those species with relatively lower pollen-attraction costs (trade-off 3 in Figure 1b). (Note that *choosiness* as defined here encompasses a number of processes that occur between ovule maturation and the onset of zygote provisioning, including pollen-limitation, pollen-ovule incompatibility, and selective embryo abortion. However among these processes, it is selective abortion that is expected to be stronger in species with a relatively higher ovule count, i.e. parental optimists.)

### The three trade-offs combine to form a single reproductive strategy continuum

The count-size trade-offs and parental optimist-parental pessimist trade-off emerge from different bodies of literature, but by extending them to consider total reproductive investment and counts of parts at two key times in a plant’s reproductive cycle, it becomes apparent that they represent the same reproductive strategy continuum and together predict a syndrome of traits associated with large-seeded (depicted in Figure 1c) versus small-seeded species. Consider a large-seeded species, one lying at the low seed count-high reproductive costs end of trade-off 1 (Figure 1b). Such species will align with the high choosiness-low relative pollen attraction costs end of trade-off 3, for species with high reproductive costs will be most selective about which embryos to provision (connection 1 in Figure 1c). A species with low seed count and high choosiness (low seedset) must as a matter of logic produce a relatively larger ovule count, aligning these species with the high ovule count-low pollen-attraction costs end of trade-off 2 (connection 2 in Figure 1c). Indeed, trade-off 3 is nearly a ratio of the two energy pool-count trade-offs: it reflects what decisions plants make after allocating energy to pollen-attraction (trade-off 2), but before beginning to allocate the provisioning component of total reproductive investment (part of trade-off 1).

In summary, at one end of the spectrum are species that produce relatively few, but large seeds, and have low seedset. These parental optimists display greater selectivity in which zygotes to provision, since they are investing more energy in each offspring and maturing fewer seeds. These species invest relatively more in seed provisioning and relatively less in pollen attraction per ovule (Figure 1c). Parental pessimists, relative to the parental optimists, have the same energy to invest in ovules or seeds, but produce relatively fewer, more costly ovules and relatively more, less costly seeds. Two previous studies have indeed observed that big-seeded species have lower seedset, also attributed to greater choosiness (Lord & Westoby 2006, 2012).

Based on these trade-offs we predict that the proportion of reproductive energy going to the different outcomes in Figure 1a will shift with seed size: 1) In large-seeded species total pre-provisioning investment will be predominately into discarded tissues, as most of the ovules produced will be shed or aborted before the onset of provisioning. 2) Once large-seeded species begin provisioning a zygote they are more likely to successfully create a viable seed, such that the proportion of total provisioning investment allocated to successful tissues versus discarded tissues should be higher in large-seeded species. 3) With increased seed size, species spend a decreasing proportion of their success costs on pollen-attraction costs, as they are expected to produce a large number of inexpensive ovules.

Overall, we ask the following questions:

1. How much do individual plants invest in different reproductive tissues and does the proportional investment differ among species?
2. Do the hypothesized trade-offs exist between pollen attraction costs and ovules available for pollination and between success costs and seed count?
3. Is there a trade-off between choosiness and pollen-attraction costs?
4. Does the proportion of energy allocated to different reproductive tissue types shift with seed size?
5. Within a species, do total accessory costs or particular accessory cost components shift with plant size, age, or reproductive investment?

The dataset we use to address these questions is, to our knowledge, the most complete dataset where plant size, vegetative investment, reproductive investment, seed investment, seed count, and seed mass were simultaneously measured across multiple species at different size and ages in a native community. In a recurrent-fire coastal heath community, we studied fourteen species differing in seed size, lifespan, and maximum height. Individuals were sampled at different ages across a fire-created chronosequence, from 3 months to 30 years. We assessed total reproductive investment every 3 weeks for a year, to determine total investment both in tissues that developed into mature seeds and in tissues that were aborted during the developmental trajectory. This detailed accounting allows us to investigate correlates of reproductive tissue pool investment across and within species.

Finally, given the complexity of measuring all the components of reproductive investment, we assess how well different surrogate measures potentially predict total reproductive investment. For this purpose we consider variables including total seed mass, total fruit mass, and total investment to the point of pollination.

## Methods

### Study system

The study was carried out in Kuring’gai National Park, just to the northeast of Sydney, Australia. The sandstone surfaces throughout the park host a coast heath community, whose dynamics have been governed by fire for at least 6000 years (Kodela & Dodson 1988). Fire regimes under traditional aboriginal management are unknown, but current New South Wales National Parks and Wildlife Service (NSW NPWS) management practises seek to achieve an average interval between 7–30 years to maintain the current floristic diversity (NSW Office of the Environment 2006). The community includes perennial species that re-sprout following fire and also obligate seeders, species that are killed by fire and re-establish from seed. The obligate seeders included in this study germinate within a year of the fire and often after the next rain. Since the fire history of the park is well documented, the age of obligate seeders at a site can be estimated. In total, we selected 14 obligate-seeder, woody perennials that are common in the community, with asymptotic heights ranging from 0.5 m – 5 m. They were *Banksia ericifolia* (Proteaceae), *Boronia ledifolia* (Rutaceae), *Conospermum ericifolium* (Proteaceae), *Epacris microphylla* (Ericaceae), *Grevillea buxifolia* (Proteaceae), *Grevillea speciosa* (Proteaceae), *Hakea teretifolia* (Proteaceae), *Hemigenia purpurea* (Lamiaceae), *Leucopogon esquamatus* (Ericaceae), *Persoonia lanceolata* (Proteaceae), *Petrophile pulchella* (Proteaceae), *Phyllota phylicoides* (Fabaceae), *Pimelea linifolia* (Thymelaeaceae), *Pultenaea tuberculata* (Fabaceae). The family Myrtaceae is well represented in the community, but absent from the study, as all locally common Myrtaceae re-sprout following fire. All sites were chosen to have minimal *Eucalyptus* cover, such that *Banksia ericifolia*, *Hakea teretifolia*, and *Allocasuarina distyla* (not included in our study because it is dioecious) would be the dominant canopy species late in succession, at heights of 3–5 m.

### Field measurements

The study was conducted over a single year, with the initial plant measurements and subsequent harvest conducted during the late autumn and early winter, the period of minimal vegetative growth in this plant community. Repeat visits were made throughout the year to record reproductive activity. Individuals were sampled at different ages across a fire-created chronosequence, from 3 months to 30 years. Site ages were estimated from fire records maintained by NSW National Parks and Wildlife Service. At the conclusion of the study, the approximate ages of the individuals on the six sites were: 1.4, 2.4, 5, 7, 9 and 31 years. Plants were tagged during May-June 2012 and harvested during May-June 2013, with a given individual tagged and harvested within 2 weeks of the same calendar date. Only one species, *Persoonia lanceolata*, displayed any shoot extension during these months. These months are similarly a period of minimal reproductive activity – only *Banksia ericifolia, Grevillea speciosa*, and (occasionally) *Hemigenia purpurea* flowered during this period – although a number of species had immature fruit from the previous year *(Persoonia lanceolata*) or small buds that would open in the subsequent year (*Boronia ledifolia, Conospermum ericifolium, Epacris microphylla, Grevillea buxifolia, Leucopogon esquamatus*).

Seven healthy individuals of each species were selected at each site (and thus age). At the beginning of the study year, basal diameter was recorded approximately 10 mm above the base to avoid the basal swelling. At the end of the study year, diameter was remeasured at the same location. Plants were then harvested at ground level and oven dried at 60°C for at least 1 week. Leaves and stems were separated and weighed.

Flowering parts on all individuals were recorded during repeat censuses, every four weeks during cooler months and every three weeks during spring and summer. At each census, all flowering parts were counted, including buds (by size class), flowers, young fruit (by size class), and mature fruit. For some species the size of immature and mature fruit and cones was also measured, as the final size of the structures was quite variable. The exact flowering parts considered varied considerably by species due to their diverse floral structures. Flowcharts detailing what flower parts were included for each species are provided in the Supplementary Material Tables S5-S18 and Figures S1-S14. The Supplementary Material also includes a table that indicates how each flowering part was counted and/or measured for each species. Each of the flower parts was independently collected from multiple untagged individuals in the community to determine its dry mass.

### Calculating reproductive investment and cost components

Total reproductive effort (RE) is the sum of investment in all the different flowering parts during the year, tabulated on a dry mass basis. For each species, reproductive parts were designated as either forming up to the point of pollination (pollen-attraction; i.e. the flower) or post-pollination and were summed into one of the two respective investment pools. For floral parts that were present at the time of pollination and continued to develop into either the seed or packaging and dispersal tissues post-pollination, the fraction of the final mass present at the time of pollination was designated part of the pollen-attraction investment and the remaining fraction as part of the packaging and dispersal investment. All calculations were made on an individual basis, although the average mass of many plant parts are based on species-level measurements. These calculations yielded total pollen-attraction tissue investment and total provisioning tissue investment. Total pollen-attraction costs and total provisioning costs are calculated by dividing the respective investment values by seed count.

To calculate the three success cost components, pollen-attraction costs, packaging and dispersal costs, and seed mass, the unit mass of reproductive parts required for the successful creation and provisioning of a single propagule were summed together. For pollen-attraction tissues, unit mass was determined by dividing the mass of the part at the time of pollination by the number of ovules it supported. All calculations make the assumption that each species produces a fixed (average) number of ovules per flower, but individual-level calculations are made for cones or inflorescence stalks which support variable numbers of flowers and hence ovules. For packaging and dispersal tissues, the unit mass was calculated by dividing the mass of the part at seed maturity by the number of seeds it supported. For seed mass, we chose to designate the endosperm and embryo as the primary reproductive unit, for it provides a consistent comparison of tissue mass across species. It is hereafter referred to as *seed size*. In contrast, the propagule includes the seed coat, and additional dispersal tissues in some species, but not others. See the Supplementary Material for a depiction of the parts for each species and the number of ovules in each part.

Discarded pollen-attraction tissue costs were then determined as:

Total pollen-attraction costs – Successful pollen-attraction costs.

Discarded provisioning tissue costs were then determined by the following formula, where successful provisioning costs is the sum of seed size and successful packaging and dispersal costs:

Total provisioning costs – Successful provisioning costs.

Reproductive count values used in the manuscript are defined as follows: *Ovule count* indicates the count of all ovules initiated by the plant. *Reach flowering count* indicates the count of ovules that developed to maturity and were presented to pollinators. *Post-pollen count* indicates the count of ovules that experienced at least some provisioning and is divided into *seed count*, the count of mature seeds formed, and *post-pollen aborted count*, the count of zygotes that aborted after provisioning had commenced. All counts are for a one-year time period.

Further detailed information on the calculation of all reproductive tissues is provided in the supplementary information.

### Statistical methods

Bivariate relationships among the variables were quantified using two methods. When testing for a significant correlation between two variables we report the r^2^ and p-value of an ordinary linear regression. When testing whether the slope of a particular trade-off or relationship differs from a specified value, we report the slope of the Standardised Major Axis line fit to the data (Warton *et al.* 2006). All analyses were conducted in R 3.2.4 (R Core Team 2015) using the package ‘smatr’ for comparing slopes of SMA lines (Warton *et al.* 2012). In addition, the code replicating this analysis (and all figures) is available at https://github.com/traitecoevo/reproductive_allocation_kuringgai (doi: will be added at proof stage).

## Results

### Accessory costs and accessory cost components

Of the 599 plants included in this study, 223 individuals produced at least one seed during the year. Across these individuals, on average 97.5% of reproductive investment went to accessory tissues rather than to seeds, decreasing to 91.5% if the entire propagule mass was treated as direct investment in offspring instead of just the embryo and endosperm components. Hereafter, all results report results for the embryo and endosperm component, designating them as seed size. Across species, accessory costs ranged from a low of 95.8% for *Epacris microphylla* to a high of 99.8% for *Hakea teretifolia* (Table 1).

**Table 1.**
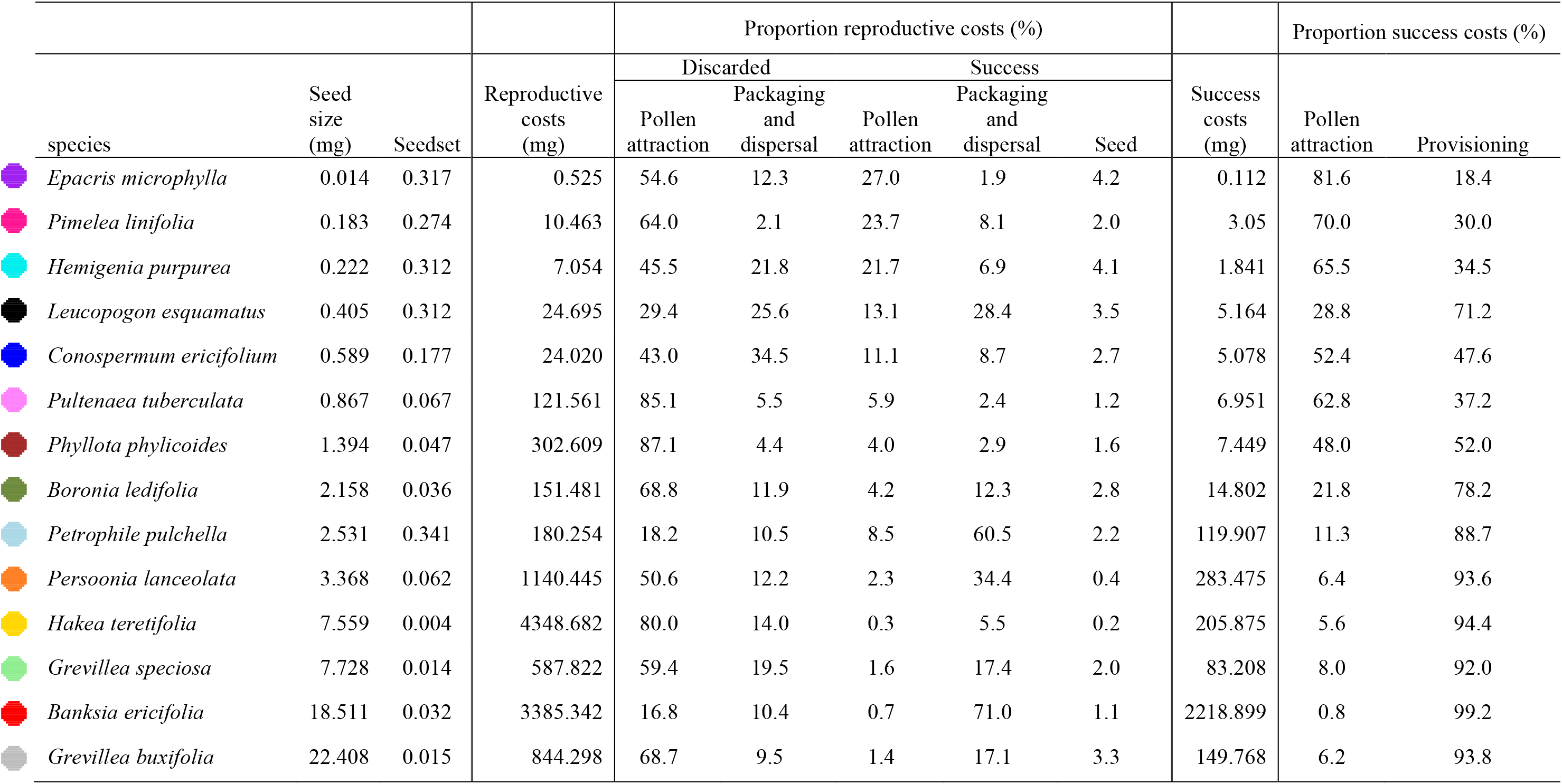
Reproductive investment data for each species. Seed size indicates the mass of the embryo and endosperm only (mg). Seedset is mature seeds per ovule initiated. Reproductive costs are the total reproductive investment per seed matured. The proportion of reproductive costs allocated to discarded tissues formed for pollen-attraction versus packaging and dispersal, successful pollen-attraction tissues, successful packaging and dispersal tissues and the seed itself are shown. Success costs are the components of total reproductive costs required for the formation of a successful seed, and are divided into two components, pollen attraction costs and provisioning costs. Note that for seed costs, the weight of the seed itself is considered part of provisioning costs. Colored dots indicate plotting colors used for each species in Figure 3.

Total reproductive costs can be divided into discarded tissue costs (the mass of all aborted and discarded parts, including mature flowers that fail to set seed) versus reproductive success costs (seed mass plus the total per ovule cost of required floral parts, both before pollination and during seed provisioning). Only the two cone-bearing species – *Banksia ericifolia* and *Petrophile pulchella* – had success costs that were higher than discarded tissue costs (Table 1). Three species – *Hakea teretifolia, Phyllota phylicoides*, and *Pultenaea tuberculata* – spent more than 90% of their reproductive investment on discarded tissues (Table 1). For most species, these discarded tissues were predominantly pre-provisioning, with aborted seeds and fruit a minor component of discarded tissue costs (Table 1). Note that fruit that abort after pollination but before the onset of visible provisioning were recorded as shed flowers, such that pollen-attraction costs (pre-provisioning) included costs associated with ovules aborted both due to lack of pollination and due to early maternal selection.

Total success costs are divided into mass of parts formed up to the point of pollination (pollen-attraction costs) versus the mass of the seed, packaging, and dispersal structures (provisioning costs). The relative size of these cost components shifted markedly across species (Table 1). Four species – *Epacris microphylla, Hemigenia purpurea, Pimelea linifolia, and Pultenaea tuberculata* – had pollen-attraction costs that were greater than 50% of total success costs, while 5 species had pollen-attraction costs that were less than 10% of total success costs (Table 1). The percentage of success costs invested in provisioning tissues (including the seed itself) ranged from a low of 18% (for *Epacris microphylla*) to a high of 99% (*Banksia ericifolia*) (Table 1). The maximum percentages of reproductive investment any species invested directly in seeds were 4.2% for *Epacris microphylla* and 4.1% for *Hemigenia purpurea*.

### Observed trade-offs

Plants produce many inexpensive ovules or proportionally fewer more expensive ovules, such that the relationship between ovule count at the time of pollination, scaled to the plant’s leaf area, versus pollen-attraction costs is highly significant and has a slope not significantly different from −1 (Figure 2a; r^2^=0.88, slope = −1.12, with 95% confidence interval [−0.90 – −1.41]). Similarly, plant produce a greater number of more expensive seeds or proportionally fewer less costly seeds, such that the relationship between seed count, scaled to the plant’s leaf area, and reproductive costs also has a slope of −1 (Figure 2a; r^2^=0.93, slope = −0.99, with 95% confidence interval [−0.84 – −1.17]).

**Figure 2.**
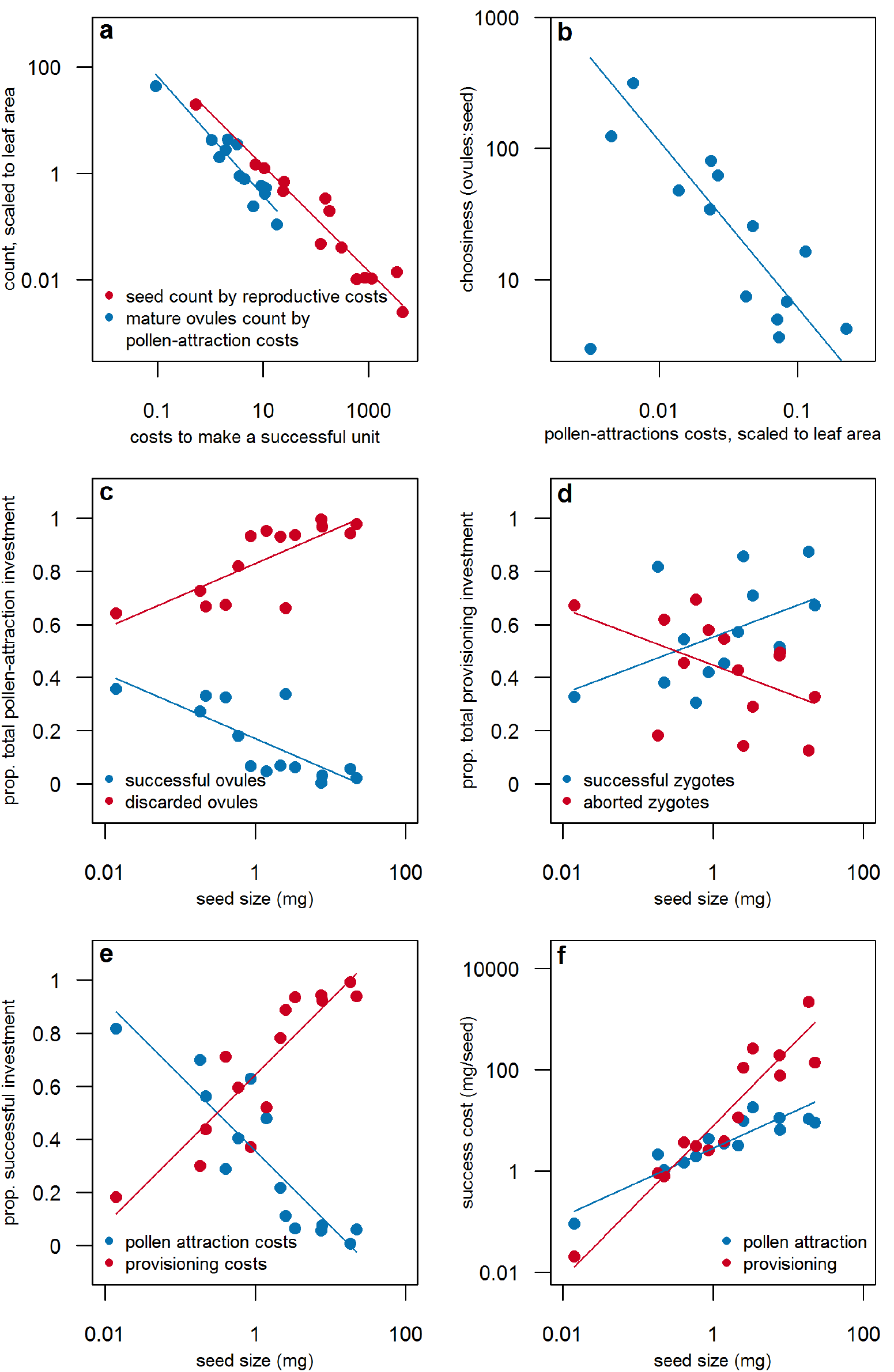
Species shift energy allocation patterns with seed size, reflecting different tissue construction costs and counts of ovules and seed produced. Each point shows average values for indivduals of a species. a) The hypothesized trade-offs between pollen-attraction costs and ovule count (r^2^=0.88) and between total success costs and seed count (r^2^=0.93) both exist. b) There also exists a trade-off between pollen-attraction costs (scaled to total leaf area) and choosiness (the ratio of mature ovules to mature seeds) (r^2^=0.76). As a result of these trade-offs, the proportion of energy invested in discarded versus successful tissues and into pollen-attraction costs versus provosioning costs shifts with seed size: c) larger seeded species invest a greater proportion of their success costs into provisioning tissues; d) larger seed species invest a greater proportion of pollen-attraction investment into discarded tissues versus successful tissues; e) there is a weak trend toward larger seeded species investing a greater proportion of their provisioning investment into successful tissues versus discarded tissues. Together, these allocation differences mean that the slope of the successful pollen-attraction costs-seed size regression is significantly lower than the slope of the successful provosioning costs-seed size regression.

There also exists a trade-off between choosiness (ovule to seed ratio, the inverse of seedset) and pollen-attraction costs, scaled to the plant’s leaf area (Figure 2b; r^2^=0.26, rising to r^2^=0.77 when *Epacris microphylla* with strangely high leaf area relative to all other metrics is removed; slope = −1.25, with 95% confidence interval [−0.91 – −1.71]). Plants which expend less of their energy budget to produce a single mature ovule, abort and discard a greater proportion of the ovules displayed to pollinators.

The values for the plotted points are listed in either Table 1 or Supplementary Material Table S1.

### Changes in relative energy investment with seed size

The strong trade-offs between the cost to produce a specific reproductive tissue and the number of units produced by the plant is manifested as shifts in the proportion of reproductive energy invested in different reproductive tissue pools across the seed size spectrum. As seed size increases, there is also a trend toward increasing expenditure on discarded pollen-attraction tissues in comparison to successful pollen-attraction tissues (Figure 2c; r^2^= 0.60, p=0.0012), reflecting the increased choosiness (decreased seed set) in larger-seeded species (r^2^ =0 .59 for the seed set-seed size regression; p = 0.0013). Increased seed size was only marginally related to a shift in the proportion of provisioning energy invested in successful versus discarded tissues, with larger-seeded species showing a slight increase in proportional investment in successful tissues (Figure 2d; r^2^= 0.24, p=0.0741). Larger-seeded species expend a greater proportion of their *success costs* on provisioning tissues versus pollen-attraction tissues in comparison to smaller-seeded species (Figure 2e; r^2^= 0.80, p<0.0001).

These shifts are also reflected in the relative slopes of the regression between seed size and provisioning costs and between seed size and pollen-attraction costs: provisioning costs show a steeper than isometric increase with seed size, while pollen-attraction costs show a less than isometric increase with seed size (Figure 2f; Table 2). The per seed matured costs of most other reproductive tissue pools show slightly steeper than isometric increases with increasing seed size, indicating the costs are relatively higher for larger-seeded species (Table 2).

**Table 2.**
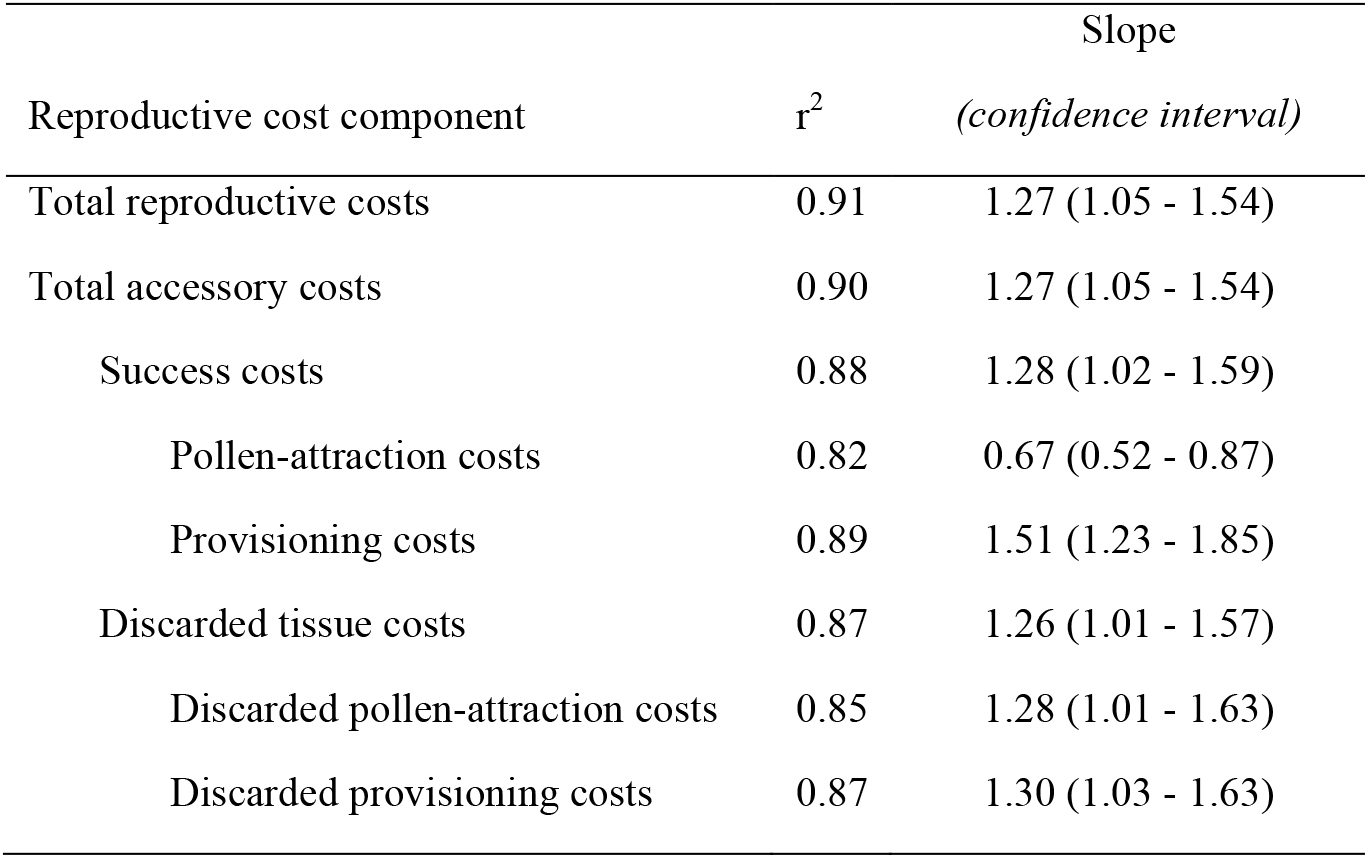
Scaling of reproductive tissue costs with seed size. All variables were showed a strong correlation with seed size (p < 0.0001). Tables show properties of SMA line fits, between different variables and seed size.

| Reproductive cost component | $r^2$ | Slope<br>(*confidence interval*) |
| --- | --- | --- |
| Total reproductive costs | 0.91 | 1.27 (1.05 - 1.54) |
| Total accessory costs | 0.90 | 1.27 (1.05 - 1.54) |
| Success costs | 0.88 | 1.28 (1.02 - 1.59) |
| Pollen-attraction costs | 0.82 | 0.67 (0.52 - 0.87) |
| Provisioning costs | 0.89 | 1.51 (1.23 - 1.85) |
| Discarded tissue costs | 0.87 | 1.26 (1.01 - 1.57) |
| Discarded pollen-attraction costs | 0.85 | 1.28 (1.01 - 1.63) |
| Discarded provisioning costs | 0.87 | 1.30 (1.03 - 1.63) |

The values for the plotted points are listed in either Table 1 or Supplementary Material Table S1.

### Shifts in accessory costs with plant size, age, or reproductive effort

None of the study species demonstrated a decrease in per seed accessory costs with increasing plant size or RE, and only one species showed a decrease in per seed accessory costs with age. With only 1/42 tests significant (Supplementary Material Table S2), this likely represents little more than chance. There are also two regressions, where accessory costs increased with plant size or age (Supplementary Material Table S2).

### Correlates with total reproductive investment

Of the 599 plants included in this study, 357 individuals produced buds and 223 individuals produced mature seeds. Even among the individuals that produced seeds, embryo and endosperm investment was only rather loosely correlated with total reproductive investment, both within and across species (Table 3, Figure 3, and Supplementary Material). All but one species showed a significant correlation between the two metrics, but only three species displayed an r^2^ above 0.80 and only eight of the species had an r^2^ above 0.70. Furthermore, the slopes and intercepts of the relationship differed across species with the result that the correlation between reproductive investment and propagule investment across individuals of all species had an r^2^ of just 0.52 (Figure 3a, Table 3). Combined these results indicate that measures of seed production alone provide poor predictors of total reproductive investment.

**Figure 3.**
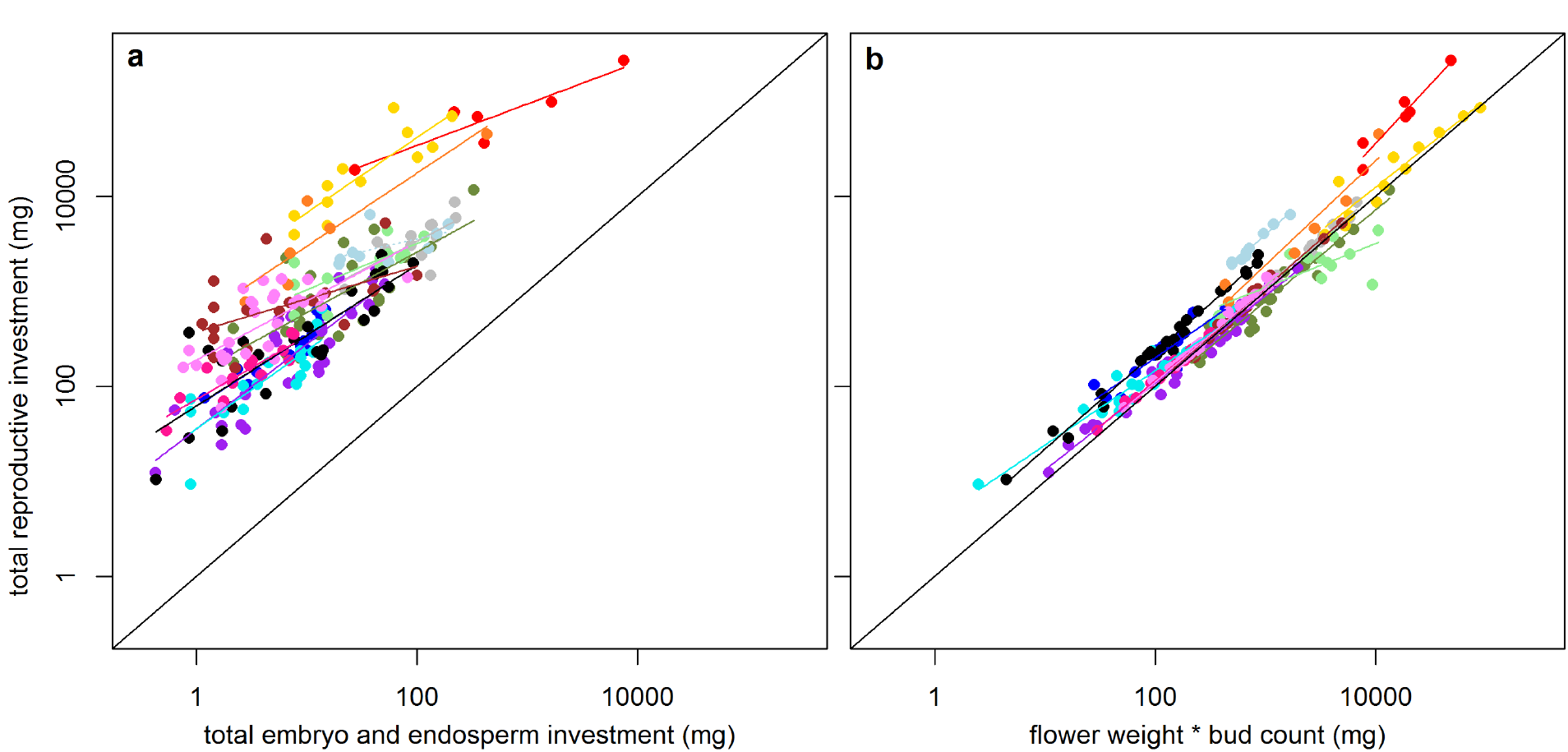
Embryo and endosperm investment is much more poorly correlated with total reproductive investment, than is a composite variable, the product of a count of the buds initiated multiplied by average flower weight. In each plot, different colored points represent the 14 study species; see Table 1 for the key. The colored lines are best fit lines through each species’ points. There are more points in panel b, as some individuals produce buds, but no seeds. In plot b, some individual’s flower weight * bud count is higher than their total reproductive investment due to a large proportion of buds aborting prior to reaching their mature flower weight. In this plot, propagule weight, the weight of the dispersed unit, not embryo and endosperm weight are used, as the purpose is to plot the commonly used currency.

**Table 3.** Correlation of different estimates of reproductive investment (and total plant weight) against total reproductive investment (mg). Regressions are done across all individuals of all 14 study species for which both reproductive investment and the *estimate* variable are greater than zero. The total cost of failed tissues or simply the energy expenditure into flowers provides the best approximation of total reproductive investment. All fits were highly significant with p < 0.0001.

| Estimate of reproductive investment | n | r<sup>2</sup> |
| --- | --- | --- |
| Total plant weight (mg) | 357 | 0.620 |
| Embryo and endosperm investment (mg) | 223 | 0.660 |
| Propagule investment (mg) | 223 | 0.525 |
| Fruit investment (mg) | 223 | 0.675 |
| Flower investment (mg) |  |  |
| (flower weight * bud count) | 223 | 0.922 |
| Successful investment (mg) |  |  |
| (success costs * seed count) | 223 | 0.728 |
| Successful pollen-attraction investment (mg) | 223 | 0.380 |
| Successful provisioning investment (mg) | 223 | 0.736 |
| Discarded tissues (mg) | 357 | 0.968 |

To assess what approximation of reproductive investment was the best alternative to measuring total reproductive investment, we regressed additional investment categories against total reproductive investment. Measures that included only investment in tissues associated with the production of mature seeds, were inferior predictors of total reproductive investment compared to measures that included investment in discarded tissues (Table 3). In particular, the correlation (r^2^) between investment in all discarded tissues versus all reproductive tissues was 0.97, while the correlation between investment in all successful tissues (success costs*seed count) versus all reproductive tissues was only 0.73. Investment in discarded tissues is a better predictor for two reasons. First, discarded tissues accounted for 73% of total reproductive investment; and second, energy investment into buds and flowers was more predictable, while further filtering processes occurred before buds become mature seeds. A composite metric, the count of buds initiated * average flower mass, when regressed against total reproductive investment, had an r^2^ of 0.92, making it nearly as strong a predictor of total reproductive investment as discarded tissue investment. Twelve of the species had the same slope for the relationship and eleven of the species had the same intercept for the relationship as the all-individuals regression (Supplementary Material S3).

## Discussion

There were four key outcomes from this study. First, we observed that plants of the14 long-lived perennial species studied expended a very large proportion of reproductive energy on accessory costs. Investment in seed dry mass represented a quite modest proportion of total reproductive investment (RE) for the 14 perennial species included in this study, with just 0.2–4% of RE going to seeds versus other reproductive tissues (Table 1). Even the individual with the lowest accessory costs invested just 9.5% of its RE into the seed itself. Second, we observed a trade-offs between ovule count and pollen-attraction costs and between seed count and total reproductive costs. The trade-offs indicate there exists a fixed pool of energy to invest and species differ in the relative cost of a part versus the number of parts they can produce. We also observed a trade-off between choosiness, the inverse of seedset, and pollen-attraction costs: species that expend less energy to produce an ovule produce a greater excess of ovules. These are species at the *parental optimist* end of the optimist-pessimist spectrum, which have proportionally costlier provisioning tissues relative to pollen attraction tissues (Rosenheim *et al.* 2014). In combination, these trade-offs lead to systematic differences in the way reproductive energy is allocated across species, resulting in a syndrome of reproductive traits values observed for large-seeded versus small-seeded species, our third outcome. The *parental optimists* were, as predicted, the large-seeded species: part of the big seed-size, low seedset strategy is to invest proportionally less in flower construction to the point of pollination and proportionally more in provisioning tissues. The fourth major result was that for perennial species with low seedset, total reproductive investment was best predicted by energy expenditure in buds and flowers, not by investment in seeds or fruit.

### Accessory costs are large

All species in this study allocated an enormous proportion of RE to accessory costs, both accessory success costs and discarded tissues (Figure 1a, Table 1). Many estimates of plant energy investment in reproduction do not account for total accessory costs, leading to potentially misleading results (reviewed in Obeso 2002; Lord & Westoby 2006; Rosenheim et al. 2014; Wenk & Falster 2015). For example, studies seeking to estimate the cost of reproduction may reach erroneous conclusions if they record only shifts in seed production year upon year, ignoring investment in accessory tissues (Obeso 2002). Reproductive allocation, the proportion of energy spent on reproduction rather than on growing and replacing vegetative tissue (Ashman 1994; Bazzaz, Ackerly & Reekie 2000; Wenk & Falster 2015), will also be substantially underestimated, leading to overestimates of the proportion of energy (and absolute amount of energy) available for vegetative growth. The current study indicates that fair assessment of RE needs to account for all pools of accessory tissues, since both discarded tissue costs and success cost components (see Figure 1a for definitions) contributed to the high accessory costs (Table 1).

Our study species have diverse floral and fruiting structures, such that disparate tissues comprise success cost expenditures in different species (Figure 1a, Table 1). For three species *(Epacris microphylla, Hemigenia purpurea*, and *Pimelea linifolia*), the costs of producing pollen-attraction tissues (on flowers that eventually produce mature seeds) was 21–27% of total RE, while for other species it was substantially less (Table 1). The two cone-producing species, *Banksia ericifolia* and *Petrophile pulchella*, had the costliest packaging and dispersal tissues, spending 71.0% and 60.5% of total RE, respectively. Other species also had high packaging and dispersal expenditure due to structures including fleshy fruit (*Persoonia lanceolata*), woody seedpods (*Grevillea* species), and thick seedcoats (*Leucopogon esquamatus*). These are tissues that must be produced to mature each seed and their exact structures have presumably evolved to maximize seed production and survival.

Discarded tissues, those tissues associated with ovules that abort instead of developing into a mature seed, are the complement to success investment. For 12 of the 14 study species, discarded tissues accounted for more than 60% of total reproductive investment (Table 1). Only in *Banksia ericifolia* and *Petrophile pulchella*, the two species with very high energy investment in woody cones, was a smaller proportion of RE attributable to discarded tissues. The majority of discarded tissue costs was due to buds and flowers that were aborted before seed provisioning became substantial (Table 1). Indeed, a large energy investment in discarded tissues has been found for all species that display low seed or fruit sets (Stephenson 1981; Sutherland 1986; Ramirez & Berry 1997; Knight *et al.* 2005).

These high accessory costs, and in particular high discarded costs, should presumably be considered a cost of sex. That is, the only reason for incurring them would be in order to create zygote genomes that conferred superior fitness, compared to zygotes created by selfing or apomixis. Having a surplus of ovules, relative to the number of offspring that can be provisioned to maturity, allows the plant to be selective about which zygotes to mature. Explanations for the abortion of a large number of mature ovules near the time of pollination include environmental stochasticity, pollen-limitation, poor pollen-tube growth, pollen incompatibility, selective abortion, and resource limitation (Ashman *et al.* 2004; Knight *et al.* 2005; Ruane, Rotzin & Congleton 2014). Additional zygotes will be lost during the provisioning period due to factors including insect attack and poor environmental conditions.

In the following sections we explore whether the three trade-offs that are observed predict how relative investment in different accessory cost pools shifts across species.

### Count-cost and choosiness-cost trade-offs exist

The first two trade-offs identified in the introduction describe how a given pool of energy can be divided into many small units or proportionally fewer large units. Abundant theory and empirical evidence underpins the seed size-seed number trade-off (Smith & Fretwell 1974; Moles *et al.* 2004; Sadras 2007) and here we extend the theory to include two trade-offs that account for the significant accessory costs required for seed production. The first trade-off is between seed count and total reproductive costs, closely related to the well-established seed size-seed count trade-off, demonstrating that large-seeded species are those species with high overall per seed reproductive costs and low seed counts (Smith & Fretwell 1974; Rees & Westoby 1997; Moles & Westoby 2006). The second is the ovule count-pollen-attraction costs trade-off, suggesting plants have a fixed pool of energy to allocate to construct flowers to the point of pollination and this energy may be divided into fewer showier flowers or into more numerous but cheaper flowers (Rosenheim *et al.* 2014).

The third trade-off is between choosiness (inverse of seedset) and the relative cost of producing a single ovule to the point of pollination: species for whom producing an ovule is less costly tend to have lower seedset (Lord & Westoby 2006; Rosenheim *et al.* 2014). Species with low seed set are also termed parental optimists: they produce excess pollinated ovules, relative to the seeds they can provision in an average year, because they are always optimistic that the year will be better than average. Due to the large number of ovules they produce, they are selected to reduce their pollen-attraction costs (Haig & Westoby 1988; Schreiber *et al.* 2015; Rosenheim *et al.* 2015). Since these species have lower seed output, they are under stronger selection to produce seeds that will successfully establish (Lord & Westoby 2006). Simply being larger is part of their strategy (Moles & Westoby 2006), but ensuring their seeds have vigorous genotypes is another correlate of this same strategy dimension and one achieved through greater choosiness for the most vigorous embryos shortly after pollination (Westoby & Rice 1982; Willson & Burley 1983; Sutherland 1986; Guittian 1993). Having excess ovules pollinated means parental optimists can be more selective in terms of pollen receipt (Zimmerman & Pyke 1988) and which zygotes to provision (Willson & Burley 1983; Sutherland 1986; Guittian 1993).

### Coordinated shifts in reproductive energy allocation across species

Together, the three trade-offs predict a single axis of variation in reproductive strategies, showing how species exhibit coordinated shifts in resource allocation, leading to a syndrome of reproductive traits associated with large-seeded versus small-seeded species (Figure 1c). At one end of the spectrum are parental optimists, using their pool of pre-pollination energy to produce many, inexpensive ovules, but their total pool of reproductive energy to produce relatively few, costly seeds, resulting in low seedset. The parental pessimists fall on the opposite end of the spectrum. As a result, species are expected to be under strong selection to coordinate their relative investment in the different energy pools described in Figure 1a. The first and third of the predicted relative shifts in tissue investment with seed size were strongly borne out by our data, while support for the second was weaker. First, since large-seeded species had lower seedset – and in particular high ovule and embryo abortion near the point of pollination – they spent a larger proportion of their pool of energy for pollen-attraction tissues on tissues that are discarded, relative to smaller-seeded species (Figure 2c). Second, since these large-seeded species had a small proportion of ovules passing through the many filters to reach the point of provisioning and since these embryos had likely been carefully selected, the large-seeded species were expected to provision a larger proportion of the selected embryos to become mature seeds. There was only a weak trend in this direction, in part reflecting the overall high success rate of embryos once post-pollination provisioning commenced among species of all seed sizes (Figure 2d).

Third, given that large-seeded species were producing relatively many inexpensive ovules and relatively fewer expensive seeds, the proportion of *success costs* allocated to pollen-attraction materials was expected to decrease with seed size while the proportion of *success costs* allocated to provisioning materials should increase with seed size, a pattern strongly observed among the study species (Figure 2e). This represents a fundamental shift in floral construction with seed size. In relative terms, larger-seeded species were producing larger packaging and dispersal tissues, but less costly pollen-attraction materials. This is being accomplished both through a reduction in floral size and, for some plant families, an increase in the number of ovules per flower or inflorescence (Lord & Westoby 2006, 2012). This trend can also be depicted by plotting pollen-attraction costs and provisioning costs against seed size: *Pollen-attraction costs* display a less than isometric increase with increasing seed size, while *provisioning costs* display a greater than isometric increase with increasing seed size (Table 2, Figure 2f). Identical patterns have been observed in other studies (Lord & Westoby 2006, 2012). They have been attributed in part to larger seeded-species tending to have biotic dispersal agents, with animal-dispersed species allocating a greater proportion of their reproductive energy to packaging and dispersal materials (Hughes *et al.* 1994; Moles *et al.* 2005; Eriksson 2008).

In this study, total reproductive costs and accessory costs both showed a steeper than isometric increase with seed size (Table 2), indicating the proportion of reproductive energy invested in accessory tissues is higher in larger-seeded species. Our result suggests that among our study species there are (slight) additional benefits to being large-seeded that have not been explored in this study, such as higher seedling germination and success (Moles & Westoby 2006). Previous studies have not found evidence for the increase in total reproductive costs and accessory costs with increasing seed size to be other than isometric in angiosperms (Henery & Westoby 2001; Moles, Warton & Westoby 2003; Lord & Westoby 2006; Chen *et al.* 2010; Lord & Westoby 2012). Note, that in these studies, seed size was defined as the mass of the entire propagule. When we recalculate the slopes of the relationships using total propagule size, we too observe an isometric relationship between total reproductive costs or total accessory costs and propagule size (Supplementary Material Table S4.)

### Shifts in accessory costs with plant size and age

An additional motivation for this study was to determine if accessory costs shifted with plant age, size or RE. The theoretical literature suggests that for plants to increase their allocation to reproduction (versus growth) as they grow and age, plants must realize some compounding benefit (Myers & Doyle 1983; Sibly, Calow & Nichols 1985; Reekie & Bazzaz 1987a; Kozlowski 1992). Increasing mortality with age has the effect of decreasing future reproductive value and selecting for increased current RE in older plants. If accessory costs declined with RE, making seed production more efficient, then plants should be selected to have fewer, larger reproductive episodes (Kelly 1994; Kelly & Sork 2002) or to delay reproduction until they are larger and can invest more energy in reproduction (Cole 1954; Wenk & Falster 2015). Such a pattern was not observed in this dataset. Across individuals within a species, total accessory costs and accessory cost components barely shifted with plant size, age, or total reproductive investment (Supplementary Material S2). The consistent lack of shift in per seed accessory costs (or seedset, data not shown) with RE (or bud count, data not shown) surprised us. There is a large literature on expected and observed trends in pollination and seedset with the size of the floral display, showing varied patterns (e.g. Primack 1987; Klinkhamer, de Jong & de Bruyn 1989; Ohara & Higashi 1994; Goulson et al. 1998), but the literature had not led us to expect a flat relationship for all 14 species (Supplementary Material S2). For many of the species studied here sample sizes were large and we sampled across their entire age range. We believe that if a shift in accessory costs (or accessory cost components) existed with plant size, age, or RE for these species it should have been detected in this data.

### Estimating reproductive effort

Realistic estimates of RE are essential for many research questions, for example plant functional growth models require estimates of the proportion of photosynthetic energy that is allocated to growth versus reproduction (Fisher *et al.* 2010; Falster *et al.* 2011; Scheiter, Langan & Higgins 2013), while demographic models may need estimates of seed production for a given RE (Garcia & Ehrlen 2002; Miller *et al.* 2012). The current study, along with others, has shown that plants are allocating energy to many different reproductive tissues, with a notably small proportion going to seeds. However, the detailed measurements required to account for all reproductive energy expenditure are not practical for many research projects and pointing researchers to the best rapidly-obtainable estimates of total RE would be beneficial to many.

At the individual level, embryo and endosperm investment, propagule investment, and fruit investment were relatively poor predictors of RE (Table 3). Even within species, knowing seed investment provided only a mediocre estimate for total RE, with only 8 of the 14 species having an r^2^>0.70 and one species not even displaying a significant correlation across individuals (Supplementary Material S3). In contrast total investment in discarded tissues (primarily representing investment in aborted flowers and buds), and our artificial composite measure “total bud count * average flower mass at the time of pollination”, provided strong estimates of total RE (r^2^=0.96 and r^2^=0.92 respectively for regressions across all individuals; Table 3). While total discarded tissue investment is not a “quick measure”, requiring repeat visits to the field and tedious accounting, the composite measure would work well for species where most of their buds and flowers are visible at a single point in time. Doing a single bud count and determining flower weight for the species would be a manageable prospect and give you a quite accurate estimate of total RE. This composite metric has the merits that it would be relatively easy to measure on large numbers of plants and that it effectively combines both the within and across species variation (Figure 3b, Table 3, Supplementary Material S3).

Conversely, these results demonstrate that if your research question requires seed investment or seed count as an output, estimates of RE will not accurately predict seed production. Instead, and in contrast to many herbaceous species (Shipley & Dion 1992), for perennial species with relatively low seedset, seed count or seed investment must be determined for each individual.

The explanation for the poor correlation between seed investment and RE is clear: most of these species have relatively low seedset (Table 1) and moreover, seed set is quite variable across individuals at a single site (Figure 3). The unpredictability of seedset and overall low seedset means that investment in seeds, at the individual level, cannot be predicted by any easy-to-measure metrics. Many stochastic processes, from pollinator activity to pollen compatibility to resource availability lie between bud production and seed production (Herrera *et al.* 1998; Wesselingh 2007; Gómez 2008). These processes lead to both individual and inter-annual variation in seed production (Copland & Whelan 1989; Mitchell 1997; Herrera *et al.* 1998).

### Methodological considerations

To reach meaningful conclusions about trade-offs between reproductive costs, counts, and seedset, accurate measurements of total reproductive investment are essential. Our accounting scheme is very detailed, but of course imperfect. The largest source of error is that we have not measured nectar production. Some of these species are known to produce abundant nectar, particularly *Banksia ericifolia, Hakea teretifolia* and both *Grevillea* species (Pyke 1983; Pyke, O’Connor & Recher 1993; Lloyd, Ayre & Whelan 2002). Very rough back-of-the-envelope calculations, based on studies of closely related species in nearby communities, indicate nectar production increases total reproductive investment by ~20% for *Grevillea speciosa*, 10% for *Hakea teretifolia*, and well under 5% for *Grevillea buxifolia and Banksia ericifolia*. Accounting for nectar production in our study would have the effect of increasing pollen-attraction costs (both successful and discarded) relative to provisioning costs (Pyke 1983; Pyke *et al.* 1993; Lloyd *et al.* 2002).

Are dry masses the best measures of expenditure, especially in a community growing on soils known to be very low in P (Beadle 1968)? Previous studies indicate that using the concentration of a limiting mineral nutrient to calculate nutrient allocations may be a better measure of a plant’s allocation choices (Reekie & Bazzaz 1987b; Ashman 1994; Rosenheim *et al.* 2014), but also that all currencies yield similar results. For example, nectar production, in comparison to reproductive tissues such as seeds, might seem relatively less expensive, in units of P than in units of dry weight or energy, potentially relevant for a community growing on low P soils. This is a direction for future investigations.

A persistent issue in assessing reproductive costs is that some green reproductive tissues are known to photosynthesize (Cohen 1976; Reekie & Bazzaz 1987a; Wesselingh 2007). It can be argued that their dry mass is not a fair measure of cost, with some of it being paid back from their own photosynthesis. Against this, it can be argued that all the plant’s photosynthesis should be considered a common pool of resource, and dry mass of different parts fairly reflects the relative allocation to different activities and tissue functions. We have adopted this second view.

This dataset does not address other known factors that may contribute to low seedset in this system, including pollen-limitation (Burd 2008, 2016) and environmental stochasticity. Insufficient pollen receipt may certainly be contributing to the patterns observed, but given recent theoretical treatments that suggest pollen-limitation should be more severe among parental-pessimists (Rosenheim *et al.* 2014, 2016), it is unlikely the observed trend of lower seedset among the parental-optimists is primarily attributable to pollen-limitation. Environmental stochasticity, both in terms of pollen receipt and resources to provision embryos, also selects for overproduction of embryos in parental optimists (Haig & Westoby 1988; Rosenheim *et al.* 2014). Parental optimists are so-named because they are optimistic about the number of ovules they will be able to mature and therefore produce additional ovules that can be matured when sufficient resources are available (Mock & Forbes 1995; Burd *et al.* 2009; Schreiber *et al.* 2015; Rosenheim *et al.* 2015).

## Conclusions

In summary, the correlations observed in our study indicate that seed size, ovule production versus seed production, and the magnitude of specific reproductive tissue pools are coordinated across species. While a plant’s accessory costs may be startlingly large at first glance, allocation of energy to different tissues is expected to represent an evolved strategy to maximize fitness. Identifying trade-offs between specific energy allocation choices – and then determining that energy allocation within this community matches the predicted patterns – provides a framework for understanding coordinated responses for seed size, seedset, and allocation to pollen-attraction versus seed provisioning tissues. Just as species have long been shown to follow a seed size-seed number trade-off, so do all species have the same amount of energy (relative to their leaf area) to invest in ovules, leading to a trade-off between the cost of pollen-attraction tissues and ovule count. Large-seeded, low seedset species have proportionally less costly pollen-attraction tissues and on average produce a proportionally larger excess of ovules relative to the seeds they are able to provision.

